# Effects of estrogen on Survival and Neuronal Differentiation of adult human olfactory bulb neural stem Cells Transplanted into Spinal Cord Injured Rats

**DOI:** 10.1101/571950

**Authors:** S Rezk, A Althani, A Abd-Elmaksoud, M Kassab, A Farag, S Lashen, C Cenciarelli, T Caceci, HE Marei

## Abstract

In the present study we developed an excitotoxic spinal cord injury (SCI) model using kainic acid (KA) to evaluate of the therapeutic potential of human olfactory bulb neural stem cells (h-OBNSCs) for spinal cord injury (SCI). In a previous study, we assessed the therapeutic potential of these cells for SCI; all transplanted animals showed successful engraftment. These cells differentiated predominantly as astrocytes, not motor neurons, so no improvement in motor functions was detected. In the current study we used estrogen as neuroprotective therapy before transplantation of OBNSCs to preserve some of endogenous neurons and enhance the differentiation of these cells towards neurons. The present work demonstrated that the h-GFP-OBNSCs were able to survive for more than eight weeks after sub-acute transplantation into injured spinal cord. Stereological quantification of OBNSCs showed approximately a 2.38-fold increase in the initial cell population transplanted. 40.91% of OBNSCs showed differentiation along the neuronal lineages, which was the predominant fate of these cells. 36.36% of the cells differentiated into mature astrocytes; meanwhile 22.73% of the cells differentiated into oligodendrocytes. Improvement in motor functions was also detected after cell transplantation.

## Introduction

Spinal cord injury (SCI) is any damage to the spinal cord leading to a change, either temporary or permanent, in the normal motor, sensory and autonomic function of the cord. Globally over 20 million individuals suffer from paralysis caused by SCI ***(Lee et al., 2014).*** The number of SCI patients is continually increasing, and each year, an estimated 180,000 individuals over the world get a new injury ***(Lee et al., 2014).***

The severity of injury and its location on the spinal cord determine the symptoms of SCI. Symptoms may vary from partial to complete loss of sensory and motor function of the arms, legs and/or body. Disturbance of breathing, heart rate, blood pressure, bowel or bladder controls are considered the most severe forms of the injury ***(Yılmaz et al. 2014)***. The main causes of SCI are motor vehicle accidents, acts of violence, sports injuries, and some diseases such as osteoporosis, arthritis and cancer of the spinal cord ***(Bellucci et al., 2015 and McCaughey et al., 2016).***

The pathophysiology of spinal cord injury is best described as bi-phasic, involving both a primary (cause of injury) and secondary phase. The primary phase of injury followed by rapid and progressive secondary injury events include physiological, anatomical and neurochemical changes leading to permanent neuronal cell death and glial scar and syrinx formation ***(Choo et al., 2007).***

Various models of SCI are based on surgical methods, which are determined by the aims of the particular research ***(Akhtar et al., 2008)***. In this study the KA was used to induce an excitotoxic SCI model. The induction of KA as a very effective excitotoxin was first done by ***Olney, (1974)***. KA has been widely utilized as a specific agonist for ionotropic glutamate receptors (iGluRs) to mimic the effect of glutamate in neurodegenerative models. It is 30-fold more potent in neurotoxicity than glutamate ***(Bleakman and Lodge 1998).*** The pathological changes following KA injection are attributed to the presence of kinate and glutamate receptors in rat spinal cord (***Monaghan et al., 1983).*** KA exerts its action by binding to its receptors and stimulates glutamate receptors to release endogenous glutamate. The over stimulation of glutamate receptors produces (1) neuronal membrane depolarization, (2) rapid Ca^2+^ influx, and (3) Ca^2+^-dependent enzyme activation and reactive oxygen species (ROS) generation. Excessive Ca2+ and ROS cause mitochondrial dysfunction, DNA fragmentation and nuclear condensation. This subsequently stimulates excitotoxic neuronal death cascade events associated with activation of caspase, astrocytes and microglial cells ***(Chen et al., 2005 and Ravizza et al., 2005)***.

***McDonald et al., (1999)*** indicated that not only the immature spinal cord environment, but also the adult one may permit neuronal differentiation under certain conditions. They were first to achieve substantial neuronal differentiation of embryonic NSCs in the adult spinal cord. Several types of stem cells have been investigated in SCI. Each cell type has a different therapeutic potential, varying according to their cellular behavior, post-implantation survival, proliferation and specific differentiation. Although each cell type has a specific mechanism for the treatment of neurodegenerative disease, they all fall into broad categories: axonal remylination, lost neuron regeneration, immune modulation, extracellular environment modification or a combination thereof ***(Vawda et al., 2012).***

Human adult OBNSCs are an ideal autologous source and interesting tool for cell-based therapy of neurodegenerative disorders. They (1) possess all of the benefits of histocompatibility, biosafety, and neurophysiological efficiency; and (2) avoid political and moral questions connected with the use of human embryos or heterologous material ***(Casalbore et al., 2009)***. ***Stefano et al. (2000)*** demonstrated that human OBNSCs under ideal conditions can proliferate like embryonic stem cells and are able to differentiate into neurons, astrocytes and oligodendrocytes.

SCI is a complex event and many obstacles persist after injury that would not be fully overcome by stem cell transplantation alone. Thus combining complementary strategies would be required to advance stem cells-based treatments to the clinical stage ***(Teng et al., 2002)***. In the current study, to further enhance the survival, proliferation and migration of transplanted adult OBNSCs in the hostile microenvironment of SCI, we treated the animals with estrogen as neuroprotective, immunomodulatory and anti-inflammatory agent ***(Siriphorn et al., 2012 and samantaray et al., 2016.)*** This was done rapidly after KA injury to prevent the acceleration of primary and secondary tissue damage and to prevent microglial invasion to the injured cord.

Estrogen treatment has been potent in various models of neurodegenerative disease, including Parkinson’s disease, multiple sclerosis (MS), spinal cord injury, cerebellar ataxia, epilepsy and some models of Alzheimer’s and stroke disease (***Leranth et al., 2000***; ***Sierra et al., 2003; Sribnick et al., 2003 and Heikkinen et al., 2004***). Neuroprotection by estrogen may be attributed to many features of this hormone ***(Sribnick et al., 2003).*** Estrogen is both a potent anti-inflammatory ***(Dimayuga et al., 2005)*** and anti-oxidant agent ***(Moosmann and Behl, 1999).*** In vitro and in vivo studies show that estrogen prevents Ca^2+^ influx ***(Nilsen et al., 2002)***, decreases the calpain activity ***(Sribnick et al., 2004)*, and** decreases proteolytic and apoptotic markers ***(Linford and Dorsa 2002; Sribnick et al. 2007b).*** The attenuation of these parameters with estrogen therapy following SCI is important and essential for neurological recovery.

## Materials & Methods

### Animals

Twenty–eight male rats (weighing 250: 300 g) were housed in autoclaved plastic cages under a temperature-controlled environment of 24 ± 1°c with access to rat-chow and water ad libitum and a 12-hour light-dark cycle. The rats were acclimated to the environment for 7 days prior to the experiment. We followed the procedures adopted by the relevant IACUC at Mansoura University, Egypt.

### Experimental Design

The animals were randomly divided into five groups: The normal control group: (n=2) was left under observation without treatment, the sham control group: (n=4) was bilaterally injected intraspinally with cereprospinal fluid (CSF). The Excitotoxic lesion group: (n=6) was bilaterally intra spinal injected with K.A (1.5 μL of 2.5 Mm), the KA+ Estrogen group: (n=6) given estrogen after KA excitotoxic injury (at 15 min and 24 h after KA injury, 4 mg/kg BW estrogen was given intravenously in tail-vein). These initial doses were followed by intraperitoneal injection of five daily doses of 2 mg/kg BW, and the treated group: (n=6) given estrogen as the fourth group followed by transplantation of adult human olfactory bulb neural stem cells (h-OBNSCs) 7 days after excitotoxic injury.

### Animal Surgery and KA injection

The rats were anaesthetized by an intraperitoneal injection of Ketamine (60 mg/kg body weight) and Xylazine (20 mg/kg body weight) mixture. With electric clippers, the dorsal area of the rat was shaved from the lower back to the neck, an area extending 2 cm bilaterally from the midline. Animals were placed in a stereotaxic frame with a vertebral clamp, and the vertebral column was immobilized. An incision was made between thoracic-13 (T-13) and lumbar-3 (L-3) vertebrae according to the method of **(*Braga-Silva et al., 2007*)**. The KA was injected bilaterally and spinally via a stereotaxic apparatus similar to that used by **Yezierski et al. (1993)**. 1.5 μL of 2.5 mM K.A (dissolved in 0.9% saline) was injected using a 10-μL Hamilton syringe. Injections were made at 1.6 mm from dorsal surface and approximately 0.5 mm from the mid line ***(David et al., 1999).*** The needle was held for 2 minutes to prevent oozing of the KA solution. Muscles and the overlying skin were closed after KA microinjection. Each rat was singly housed and supplied with adequate food and clean water. Prophylactic analgesics and antibiotics were not used, to prevent any possible interaction with the experimental therapy.

### Behavioral Assessment

#### The Basso, Beattie and Bresnahan (BBB) score

The hind limb locomotor function of each animal was evaluated using a 21-point scale ***(Basso et al. 1995).* A** Zero BBB score was given when there was no movement of the hind limbs, while a score of 21 was given when trunk stability, plantar stepping, toe clearance, an erect tail and coordinated limb movement was detected (***Basso et al. 1995).*** The tests were done by two independent blinded examiners. Motor function of hind limbs of each rat was assessed for 4 minutes weekly till the end of the study. The final average score of the rats in all groups was then compared.

#### Human Olfactory Bulb NSC Isolation and Culturing

This step was described with details in our previous study published as **Marei et al., (2016)**.

#### Transfection and Infection

Human embryonic kidney (HEK)-293T cells in log-phase growth were rapidly transfected by using the standard Lipofect Amine reagent **(Invitrogen)**, with GFP-Vector plus helper plasmids to develop virions ***(Cenciarelli et al., 2006).*** Two days after cell transfection media containing virions was collected and transferred directly onto OBNSCs. Lentiviral infection was performed in the presence of polybrene solution (8 mg/ml) **(Sigma-Aldrich)**. During the OBNS/PC selection and maintenance time, the antibiotic G418 (Euroclone) was added to the cells at 400mg/ml.

#### OBNSCs Transplantation

Animals in the treated group received cyclosporine (10 mg/kg/daily, s.c) one day prior to transplantation till the end of the study. Transplantation surgeries occurred at 7 days after KA excitotoxic injury. Immediately before transplantation, cell viability was assessed by the trypan blue exclusion test (only populations with a 95 % viability were transplanted) and the total number of cells was measured using a hemocytometer. After that they were suspended at 40,000cells/μL in artificial CSF (Sigma-Aldrich), and were kept on ice during transplantation surgery. Rats were re-anesthetized as before and the site of surgery was re-opened. Each rat was stereotaxically and bilaterally injected with 4 μL (2 μL for each side) of cell suspension (160,000 cells) at a depth of 1.5mm, 4mm cranial to KA injected site. The needle was removed after 5 min to avoid cell suspension oozing. The muscle and skin were sutured again.

#### Sample collection

The animal samples in time course experiments are listed in **Table (1).** The spinal cord was collected after removal of vertebrae. Some samples were preserved in liquid nitrogen for PCR studies, and the others were fixed in 10% neutral buffered formaldehyde and were processed for paraffin sectioning.

**Table (1):** The animal samples in time course experiment.

| **Animal samples** | Control group |  | Sham control group | KA excitotoxic group |  | Estrogen treated group |  | Estrogen + OBNSCs group |
| --- | --- | --- | --- | --- | --- | --- | --- | --- |
|  | PCR | Paraffin sections | Paraffin sections | PCR | Paraffin sections | PCR | Paraffin sections | Paraffin sections |
| **1<sup>st</sup> day** |  | - | 2 | 2 | 2 | 2 | 2 | 2 |
| **1 week** | - | - | 2 | 2 | 2 | 2 | 2 | 2 |
| **6th** | 2 | 2 | 2 | 2 | 2 | 2 | 2 | 2 |

| week |  |  |  |  |  |  |  |  |
| --- | --- | --- | --- | --- | --- | --- | --- | --- |
| Total number (28 rats) | 4 rats |  | 6rats | 6rats |  | 6 rats |  | 6 rats |

#### Tissue preparation for Real time PCR

Extraction of RNA from spinal cord’s tissue was accomplished using a QIAamp RNeasy Mini kit **(Qiagen, Germany, GmbH)**. Each 100 mg of the spinal cord’s sample was disrupted and lysated in 600 μl RLT buffer comprising β-mercaptoethanol (10 μl/ml buffer) followed by homogenization of the lysate in Qiagen tissue Lyser for 2 minutes at 30 Hz speed. 70% ethanol (one volume) was pipetted and mixed with the cleared lysate. The remaining steps of total RNA isolation and purification from animal tissues were carried out using QIAamp RNeasy Mini kit protocol **(Qiagen, Germany, GmbH).** The sequences of oligonucleotides (Metabion, Germany) used for this study are listed in **Table (2).** The house keeping gene or normalized gene (B-actin) was used to estimate the relative fold change or gene expression in target gene using the 2^−∆∆Ct^ method ***(Yuan et al., 2006***).

**Table (2):**
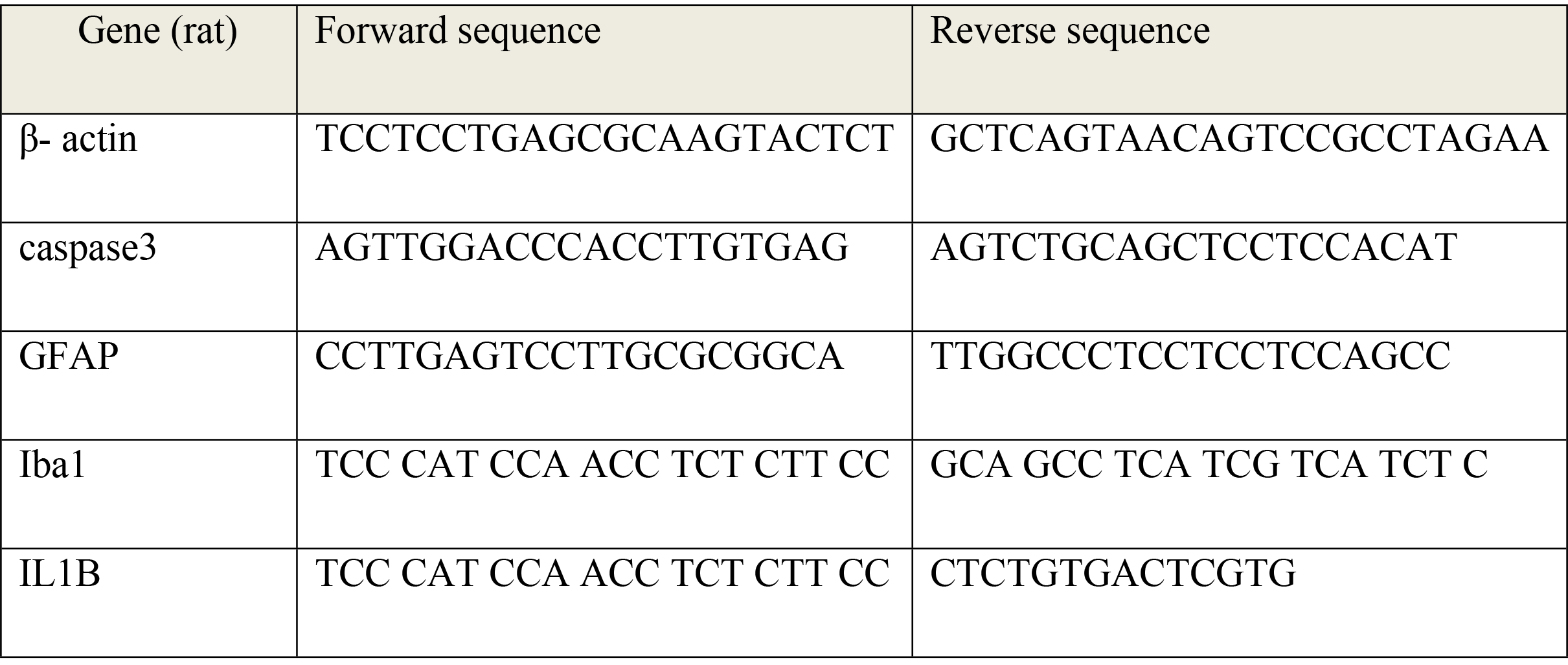
Primers sequences for qualitative rt-PCR

### Histological Analysis

#### Light microscope (LM) tissue fixation and processing

The spinal cords were removed. The injection sites were then localized under a stereo microscope to include segments from thoracic-11(T-11) to approximately lumbar-5(L-5). The cord was postfixed in the 10% neutral buffered formaldehyde for 24 h. Fixed samples were dehydrated in a graded series of ethyl alcohol, cleared in xylene and infiltrated with and embedded in paraffin. A rotatory microtome was used to cut (5 μm) transverse sections and mounted on glycerol-albumin-coated glass slides. The paraffin sections were kept in incubator at 50°C for 20 minutes then used for staining with Hematoxylin-esoin (h&E) or with Cresyl Echt Violet stains.

#### Fluorescent immunohistochemistry (IHC)

This technique was applied according to the Abcam protocol (www.abcam.com/technical.IHC-P). Briefly, transverse paraffin sections of spinal cord mounted on positive charged glass slides were processed for double immunohistochemistry with specific antibodies for human neural stem cells (Sigma-Aldrich) and examined under a fluorescence microscope. Tissue sections were deparaffinized and rehydrated using standard methods as mentioned before. The slides were immersed in a pre-boiling antigen retrieval solution (citrate buffer saline, ph. 6) for 15 minutes in a microwave (at 100°c) followed by washing with phosphate buffered saline. Tissue sections were blocked with PBS supplemented with Triton 100x (Sigma-Aldrich) and 5% normal goat serum for 1 hour at room temperature then incubated for 24 hrs at 4°C with the 1^st^ diluted primary antibody Anti-GFP (1:2000, rabbit) followed by washing with phosphate buffered saline. Sections were incubated for 1 hour in the dark at room temperature with diluted secondary antibodies [anti-Nestin (1:200, rabbit) for detection of undifferentiated NSCs; and anti-beta-tubulin III (1:100, rabbit) for detection of immature neurons, with monoclonal-anti Ng2 (1: 100) for oligodendroglia precursor cell detection; monoclonal anti-GFAP(1:300, mouse) for detection of astrocytes; and monoclonal anti O4 (1:200, mouse) for oligodendrocytes detection. This was followed by a rinse with phosphate buffered saline and application of the diluted primary antibodies: whole molecule Anti-Rabbit IgG (1:200, goat, FITC antibody, Sigma-Aldrich) then secondary antibodies (Whole molecule, Tritic conjugated–anti-rabbit IGg, 1:200, goat or Tritic conjugated–anti-mouse IGg, 1:200, goat) as mentioned.

#### Enzymatic immunohistochemistry (IHC)

This technique was applied to make stereological quantification of engrafted OBNSCs at the end of the current study. This protocol was carried out according to ***(Karen Petrosyan et al., 2002)***. Transverse paraffin sections of spinal cord (at 6^th^ week) mounted on positive charged glass slides were processed for single enzymatic immunohistochemistry with anti hGFP. Each Section was incubated with anti GFP ( the same antibody that was used in fluorescence but at **1:100** dilution) for 3hrs at room temperature, then washed. Secondary antibody (ready to use) (Biotinylated Goat Anti-Rabbit IgG, **ab64256**,) **(Abcam)** was applied to each section for 15 min at room temperature followed by washing with PBS. Freshly prepared horseradish peroxidase substrate was applied to the sections at room temperature until suitable staining developed (about 5 min); then they were rinsed with PBS and counterstained with hematoxylin for 3 minutes Slides were washed, dehydrated and examined under a light microscope.

#### Stereological Quantification

An enzyme stained section was examined under an Olympus CX41 light microscope. The approximate percentage of GFP-Positive differentiated cells (according to nuclear morphology) was assessed by a hematoxylin nuclear counterstain and GFP immunolabeling. For quantification of GFP positive cells, starting sections were selected randomly, and the four serial sections that contained the most human cells were used for fate analysis. The number of labeled cells was counted for each type (neurons or astrocytes or oligodendrocytes) using Image J Version 10.2 software with a cell counter plug-in.This number is expressed as a percentage of the total number of GFP-positive cells counted in each individual type.

#### Statistical analysis

Statistical analysis of all groups was performed using one-way ANOVA analyses P<0.05 was considered statistically significant.

## Results

### Histological and histo-pathological results

#### Control group

The spinal cord of of the rat is composed of two distinct regions; the outer white matter and the inner gray matter. The shape of gray matter was similar to that of a butterfly or H shape; within it a central canal is located. The ventral horns of H shape were broader than the dorsal horns (figure1a). The gray matter of each ventral horn is formed primarily of the large neuronal cell bodies, a mixture of glial cells, several blood capillaries, and a fine amorphous eosinophilic background called the neuropil (figure1b). The cytoplasm of neuronal cell bodies and dendrites containing basophilic granules called Nissl granules (figure1c).

**Figure1:**
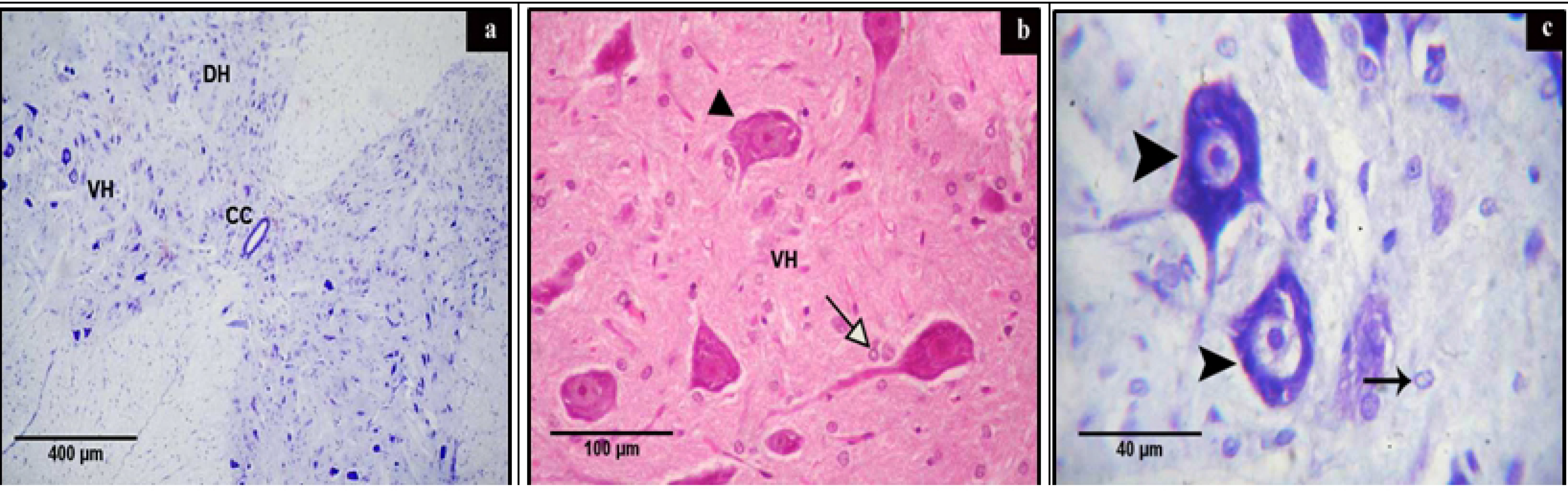
Photomicrograph of a section of rat’s spinal cord of control group showing. (a): central canal (CC) running through gray matter that was formed of dorsal horn (DH) and ventral horn (VH). Cresyl violet stain. (b): the gray matter of ventral horn containing neuronal cell body (arrow head) neuroglia cells (arrow). H&E stain. (c): gray matter of ventral horn containing astrocyte nucleus (arrow) and multipolar neurons that contain Nissl granules as basophilic mottled structures surrounding light stained nucleus with prominent nucleolus (Arrow head). Cresyl violet stains.

#### Sham control group

This sham group was examined to confirm that the degeneration that occurred in spinal cord cells was certainly a result of KA injection, not from the Hamilton needle. There was no histological alteration after CSF injection.

#### Excitotoxic lesion group

The microscopic examination of spinal cord sections revealed that KA lesions were mainly located in the ventral horn. The dorsal horn showed relative preservation of tissue architecture and its neurons were almost completely spared by K.A. injection from 1 day to eight weeks post injury. The white matter remained morphologically intact (fine meshwork-like appearance) after KA injection during the whole period of study. A variety of necrotic alterations was observed in the ventral horn of the gray matter from the first day of KA injection to the 6^th^ week after injection. The affected neurons were characterized by cell body shrinkage, intensely stained eosinophilic cytoplasm (red dead neuron) (figure2a), and loss of Nissl substance (chromatolysis) (figure 2b, c). The neuropil adjacent to the necrotic neurons was finely vacuolated or edematous with markedly astrocytic reactions(figure2a).

**Figure2:**
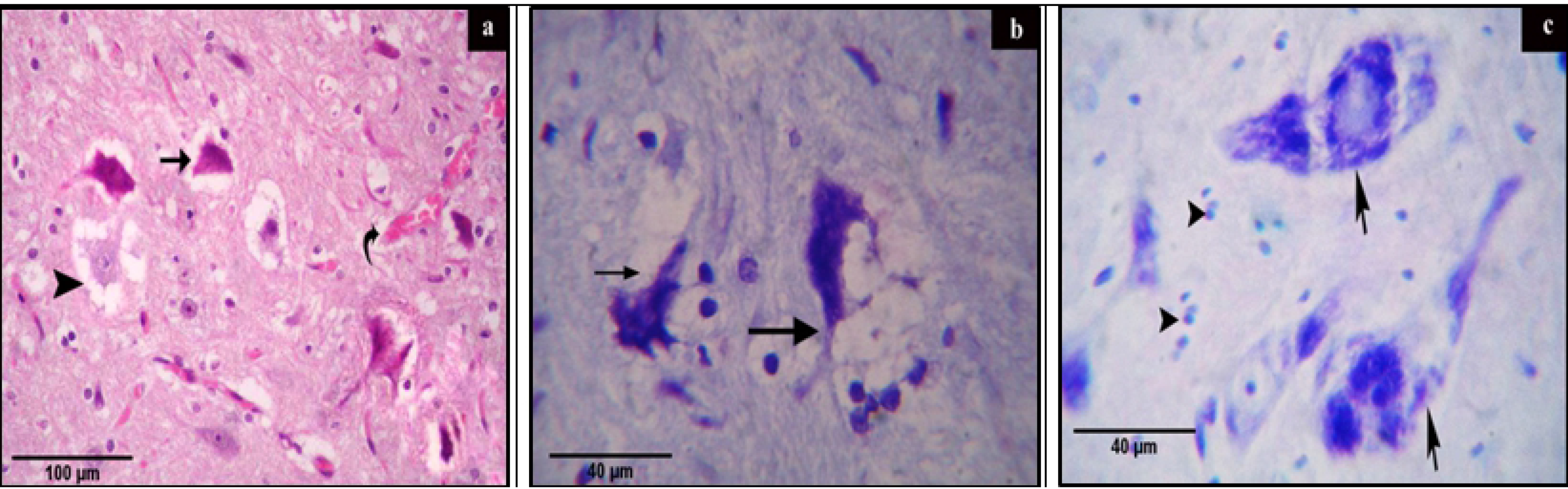
Photomicrograph of a section of rat’s spinal cord of control group showing. (a): The gray matter of ventral horn containing necrotic neuron (arrow), perineural vacuoles surrounded the remnant of necrotic neuron (arrow head) and congested blood capillary (curved arrow). H&E. (b): The gray matter of ventral horn containing debris of necrotic neurons surrounded with perinural vacuole and large number of astrocytes (arrow). Cresyl violet stain. (c): the gray matter of ventral horn containing necrotic neurons with chromatolysis (arrow) and gliosis (arrow head). Cresyl violet stain.

#### Estrogen treated group

Histopathological examination of spinal cord sections showed that the administration of estrogen rapidly after spinal cord injury dramatically improved the histological alterations following KA injury. A small number of neurons displayed signs of recovery and regeneration on the 1^st^ day of estrogen treatment (figure3a). After that the number of preserved neurons with clear integrated nissl granules was increased (figure3c). Only small numbers of necrotic neurons were still noticeable and the edematous or vacuolated neuropil wasn’t detected (figure3b

**Figure3:**
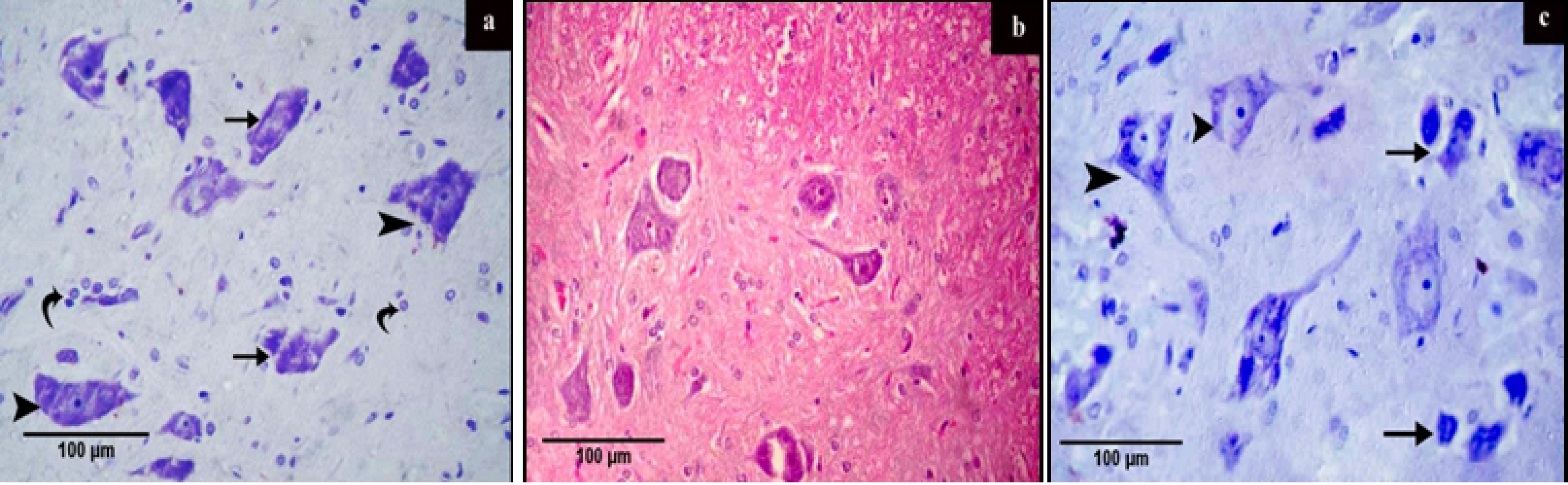
Photomicrograph of a section of rat’s spinal cord of control group showing. (a): 1day post estrogen treatment showing the gray matter of ventral horn containing necrotic (arrow), semi regenerated neurons(arrow head) and the nucleus of astrocytes(curved arrow). H&E stain (b): The gray matter of ventral horn containing apparently normal neurons with vesicular nucleus and prominent nucleolus (arrow), necrotic neuron (arrow head) and blood capillary (curved arrow). H&E stain. (c): The gray matter of ventral horn contains neurons with clear integrated Nissl granules (arrow head) and necrotic neurons (arrow). Cresyl violet stain.

**Figure 4:**
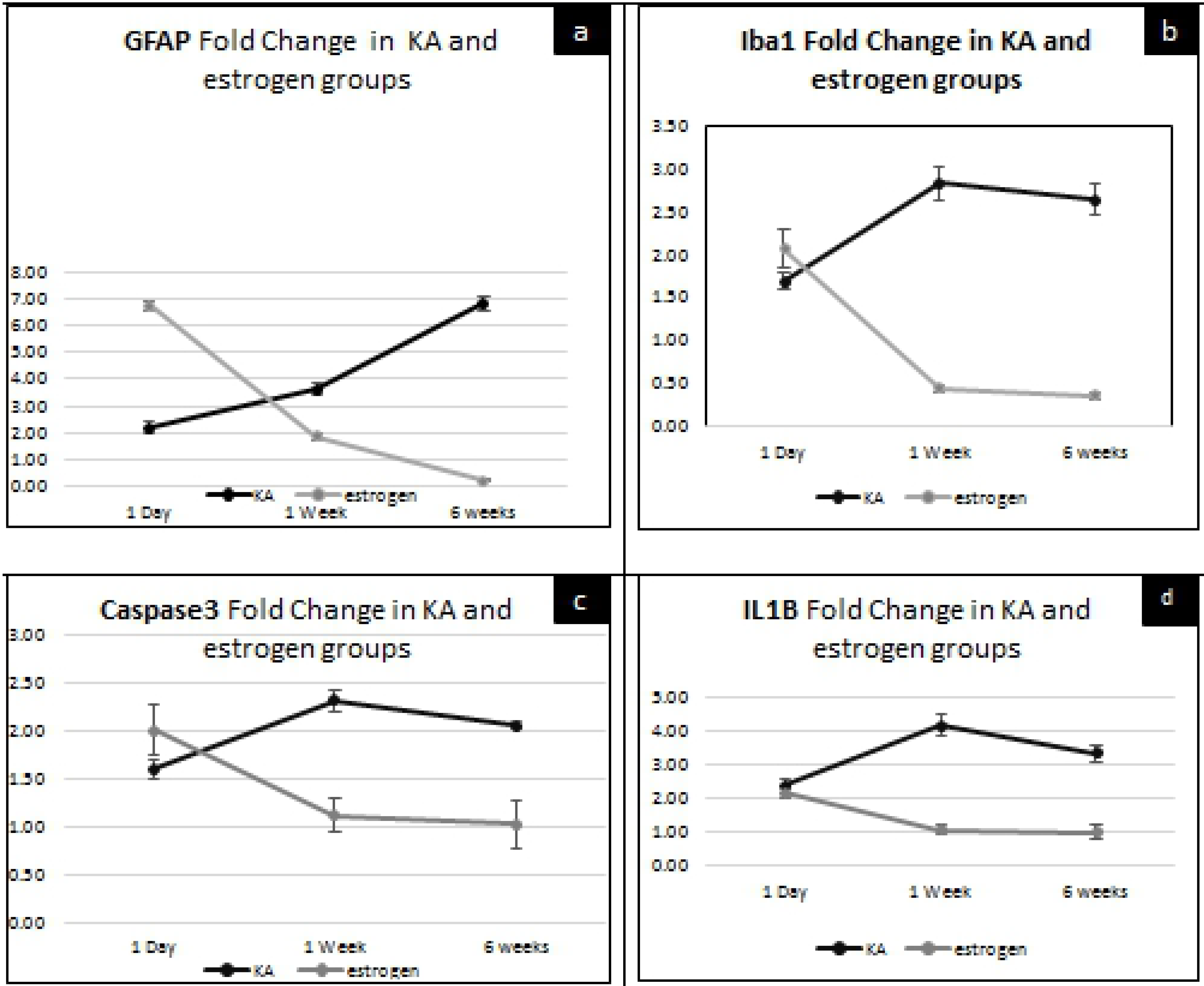
Graphs of quantative gene expressions of (a) GFAP both in KA and estrogen showing up regulation of GFAP after one day of KA injection till the end of the study, but after estrogen treatment the expression of GFAP was significantly (P-Value<0.05) decreased afer one week.(b) Iba-1in KA and estrogen groups showing up regulation of it following KA injection from 1st day till the end of the study, but after estrogen treatment this expression was significantly (P-Value<0.05) down regulation after one week till the end of the study. (c) caspase-3 both in KA and estrogen showing upregulation of caspase-3 expression after one day of KA injection with maximum increase at one month and significantly down regulation (P-Value<0.05) of it after one week of estrogen treatment. (d) IL-1β in KA and estrogen group showing significant (P-Value<0.05) up regulation of IL-1β expression after KA injection at each time point of the study and significantly decrease after three days of estrogen tratment.

### Quantitative real time PCR

In the KA group the up-regulation of caspase-3, GFAP, Iba-1 and IL-1β expressions was significantly visible **(P-Value<0.05)** on the 1^st^ day of injury (1.6 fold, 2.19, fold, 1.7 fold and 2.4 fold changes) respectively till the end of the study (2.6 fold, 6.8 fold, 2.7fold and 3.4 fold changes) respectively. In the estrogen treated group the expression of caspase-3 activity and GFAP at one day (2 fold, 6.37 fold) respectively was significantly higher than the KA group (**P-Value<0.05)**. Administration of estrogen led to obvious and significant (**P-Value<0.05)** down regulation of caspase-3 activity (1.02 fold) and GFAP expression (0.19 fold change) at 6^th^ week. The down regulation of Iba-1 (0.44 fold change) and IL-1β (1.07 fold change) after estrogen treatment was significantly detected at 1^st^ week **(P-Value<0.05)** till the end of the study (0.36 fold change for Iba-1 and 1 fold change for IL-1β).

### Immunohistochemical assessment for the engrafted h-GFP-OBNSCs

GFP was mainly used to trace the transplanted OBNSCs, and to differentiate between exogenous (our engrafted), and endogenous (non-human, non-GFP-positive) neuronal and glial elements. Immunohistochemical assessment was performed to evaluate the ability of the engrafted OBNSCs to survive, proliferate, differentiate, and restore lost or damaged neurons following the induction of the KA into spinal cord. On the 1^st^ day following engraftment, examination of double IHC sections of spinal cord (GFP positive + neuronal or glial markers) with a fluorescence microscope revealed positive immune-reactivity to nestin (figure5 a). At one week following transplantation, some of the engrafted OBNSCs showed positive immunoreactivity for GFP and GFAP (astrocytes) (figure 5 b), ß-tubulin III (immature neuron) (figure 5 c) and NG2 (immature oligodendrocytes) (figure 5 d). Later on at the 6^th^ week of transplantation, many GFP positive cells exhibited positive immune reactivity for mature neuronal markers (MAP2) (figure 5 e), the mature oligodendrocyte marker (O4) (figure 5f) and astrocytes (GFAP) (figure 5 g).

**Figure 5:**
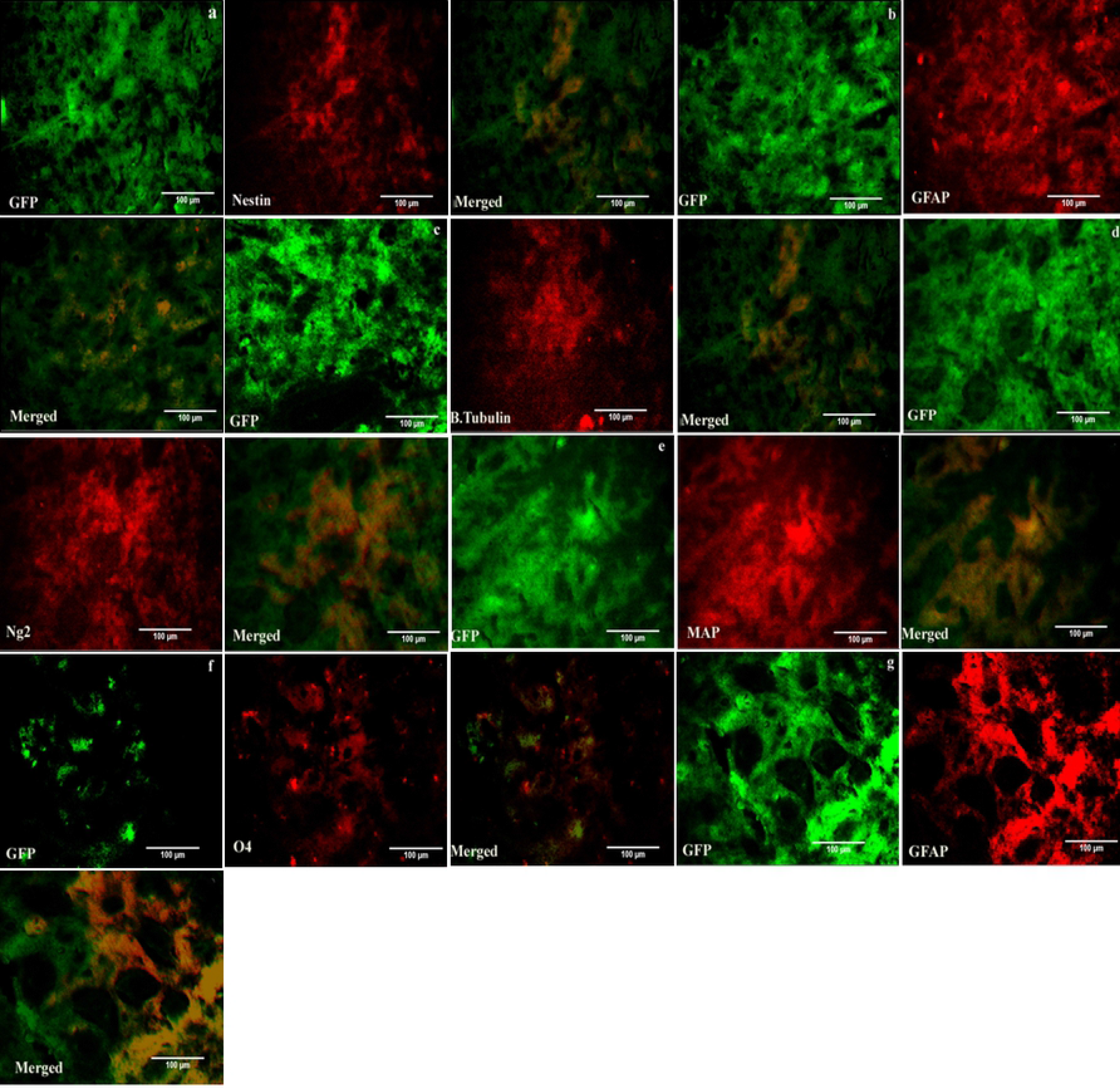
Fluorescent double-immuno histochemical staining of paraffin sections for GFP and different neural stem (Nestin), ß-tubulin III (immature neuron), NG2 (immature oligodendrocytes), MAP2 (mature neuronal), O4 (mature oligodendrocytes) and GFAP (astrocyte) markers. (a): GFP-Nestin immunoractive positive undifferentiated OBNSCs at 1st day post implantation.(b): Differentiation of OBNSCs into GFP-GFAP positive cells after one week of transplantation.(c): Differentiation of OBNSCs into GFP-ß-tubulin III after one week of transplantation.(d): Differentiation of OBNSCs into GFP-Ng2 after one week of transplantation. (E): Differentiation of OBNSCs into GFP-MAP2 6weeks post engraftment. (f): Differentiation of OBNSCs into GFP-O4 6 weeks post engraftment. (g): Differentiation of OBNSCs into GFP-GFAP 6 weeks post engraftment.

### Stereological quantification of OBNSCs

Both the number of surviving cells and the percentage of final differentiation were stereologically quantified in enzymatic immune histochemical stained sections against h-GFP. Six weeks following transplantation, GFP positive neuron-like cells, GFP positive astrocytes and oligodendrocyte-like cells with dark brown granules in their cytoplasm were detected in the different parts of gray matter. Besides these GFP-positive cells, GFP-negative (endogenous, nonhuman) cells were also observed (figures 6 a, b and c).

**Figure 6:**
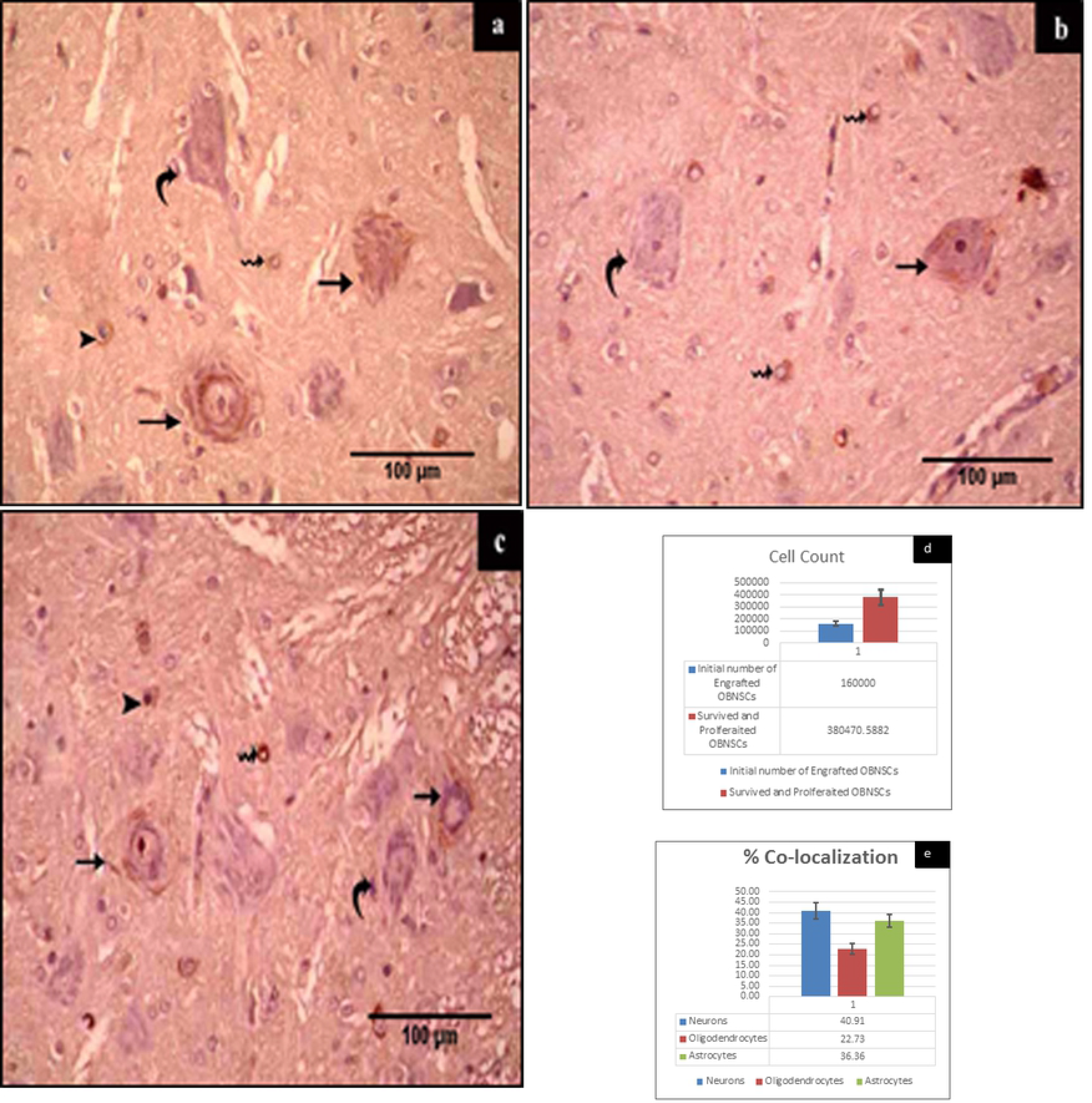
**(a, b and c):** Enzyme immunohisto-chemical staining of paraffin sections for GFP at 6th week post engraftment of h-GFP-OBNSCs showing, GFP negative neuron (curved arrow) and GFP positive cells with morphological criteria of neurons (arrow), oligodendrocytes (arrow head) and astrocytes (wavy arrows) distributed in gray matter of ventral horn. (d, e) Graphs of soteriological quantification of hOBNSCs after 6 weeks of transplantation showing (d): The initial number of transplanted cell per each rat was 160000 but after two month the average number of transplanted cell was 380407. (e): about 40.91% of transplanted cells differentiated towards neuronal linage, 36.36% towards astrocytes linage and 22.73% towards oligodendrocytes lineage.

The initial transplant contained 160,000 GFP-OBNSCs per animal; stereological estimates of h-GFP-OBNSCs (n=10) revealed an average of 380470 GFP-OBNSCs present at 6 weeks following transplantation (figure 6d). These values represent an approximately 2.38 fold increase in the initial cell population transplanted. Stereological quantification revealed 40.91% of OBNSCs showed differentiation along the neuronal lineages, which was the predominant fate of these cells. 36.36% of the cells were differentiated into mature astrocytes; meanwhile 22.73% of the cells were differentiated into oligodendrocytes (figure 6e).

### Locomotor Function Assessment (BBB score)

The data obtained from all groups were statistically analyzed by the one way ANOVA test. Assessment of hind limb motor function displayed no difference between the control and sham control groups in whole time of the study.

The average BBB score for the KA excitotoxic group, estrogen treated group, and estrogen+ OBNSCS group are collectively shown in table **(3)**

**Table(4):**
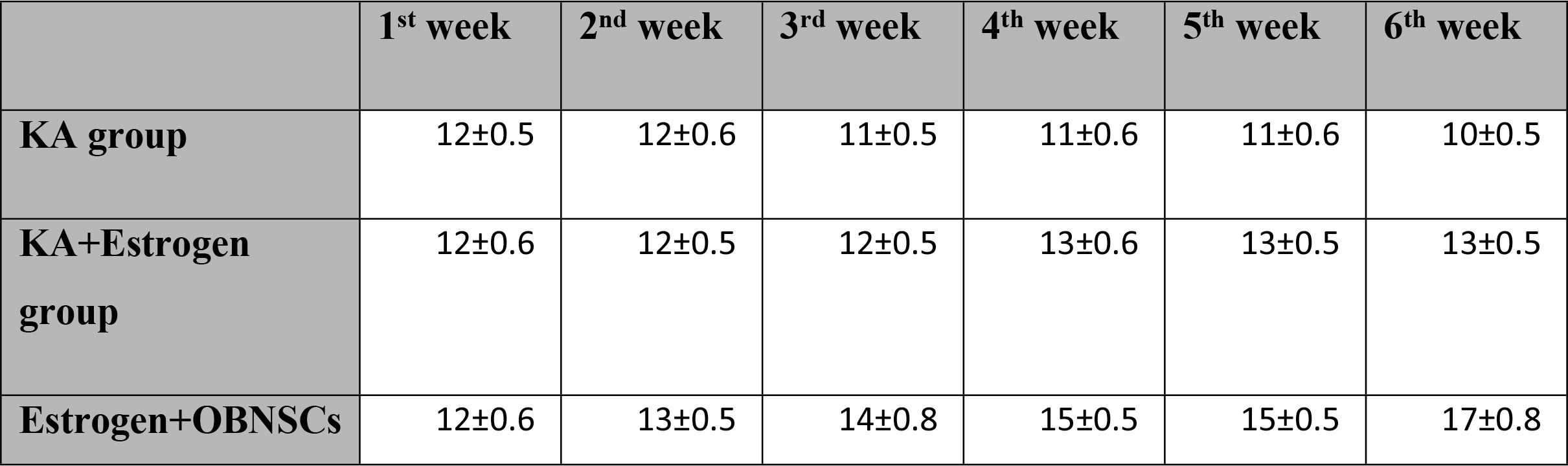
showing the BBB score for KA, estrogen and OBNSCs+estrogen groups there is no significant difference between three groups at 1^st^ week **(P-Value>0.05)**. At 2^nd^ week there is a significant improvement **(P-Value<0.05)** in OBNSCs+estrogen compared to KA and estrogen groups. After that there is a significant improvement in estrogen and OBNSCs+estrogen groups compared to KA, but this improvement was more pronounced in OBNSCs+estrogen group than estrogen group.

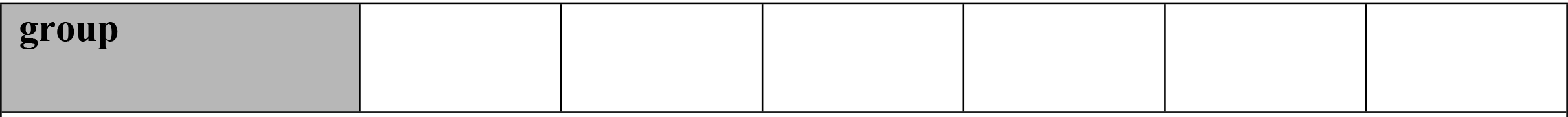

#### KA excitotoxic group

The BBB scale showed that KA excitotoxic injury caused bilateral locomotor deficits in hind limbs that persisted to the last day of the study. In comparison to control or sham control groups (BBB score 21, exhibiting normal locomotion), the average BBB score at the 1^st^ and 2^nd^ week after injury was 12 (frequent to consistent plantar step with both hind limbs and occasional coordination between fore and hind limb). From 3^rd^ to 5^th^ weeks, the animals were unable to coordinate fore and hind limbs during locomotion (average BBB score 11). At the 6th week the average BBB score was decreased again. The rats were unable to make frequent or consistent plantar steps with both hind limbs and there was no coordination between fore and hind limbs (final average BBB score 10). This indicated the success of the SCI model establishment.

#### Estrogen treated group

The average BBB score gradually increased after estrogen injection. In comparison to the KA group, the improvement in locomotor function was significant **(P-Value<0.05)** at the 3rd week of estrogen injection (average BBB score, 12). The average BBB score at the 4th week was 13, after that there was no further improvement. Functionally, this score indicated that the estrogen treated rats were frequently supporting their own body weight with plantar stepping and frequent coordination of forelimbs and hind limbs.

#### Estrogen + OBNSCs group

Assessment of hind limb motor function in the estrogen +OBNSCs group revealed consistently higher BBB scores than those of the estrogen treated group. In comparison to the KA groupthe improvement in BBB score became significant **(P-Value<0.05)** two weeks after OBNSCs transplantation (earlier than the estrogen group). By the 3rd week, the combination of estrogen and OBNSCs therapy resulted in a two-point improvement on the BBB scale compared to the estrogen group. The final average BBB score at the 6^th^ week was 17. This BBB value means that rats in this group were able to move both hind limbs with consistent plantar stepping and consistent fore/hind limb coordination. Toe clearance occurred frequently during forward limb advancement.

## Discussion

### Tissue architecture following KA injury and estrogen treatment

The results of the present work have shown that the histopathological changes following KA injection are confined to the ventral horn of spinal cord. Meanwhile, the dorsal horn and white matter showed relative preservation of tissue architecture. Our results are in agreement with the results of previous studies showing that, importantly, kinate excitotoxicity in adult rat cervical or lower thoracic and upper lumbar spinal gray matter selectively damages motor neurons responsible for forelimb or hind limb motor activity, respectively; but not long-tract motor axon transmission ***(Sun et al., 2006)***. However, another study showed severe damage of both gray matter (dorsal and ventral horn) and white matter (ventral and ventrolateral tracts) after KA injection ***(Nishida et al., 2015)***.

The integrity of tissue following estrogen therapy was significantly preserved starting from 7 days post-treatment until the end of the study. Preserving the structural integrity of the spinal cord’s injured tissue is critical for functional recovery after SCI. The neuroprotective effect of estrogen was mainly due to decrease in Caspase-3 activity, which was accompanied by a reduction of motor neuron death following injury (***Cuzzocrea et al., 2008 and Kachadroka et al., 2010)*** and inhibition of N-methyl-D-aspartic acid (NMDA) receptors (subtype of glutamate receptor) that are ordinarily activated following KA injection ***(Weaver et al., 1997).*** Our results also agree with the results of previous studies which revealed that the edema and degenerated ventral horn neurons were noticeably attenuated in estrogen treated groups compared to vehicle-treated lesion ***(Sribnick et al., 2005, 2010***, ***Siriphorn et al., 2012 and Samantaray et al., 2016)***.

### Neuronal mRNA expression following KA injury and estrogen treatment via Quantitative real-time PCR (q RT–PCR)

Examining the changes in mRNA expression after KA and estrogen injections is a powerful tool to correlate the pathophysiology of KA excitotoxicity and the histopathological findings. In the current study quantitative real-time RT–PCR with a fluorescent TaqMan probe was used to measure alterations in individual gene expression after SCI, particularly genes involved in apoptosis (caspase-3), astrocytosis (GFAP), inflammation (IL-1B) and microgliosis (Iba1). These genes were selected since they represent various cell types that are related to essential pathological processes that happen after SCI.

Caspase-3 activation has been confirmed during the process of secondary injury after SCI **(*Namura et al., 1998*).** The activation of caspase-3 after KA injury is attributed to the high level of ca+2 dependent enzymes and cytochrome c released after KA injury. Those are important activators of the caspase −3 apoptotic cascades **(*Verhagen et al., 2000*).** In the present work, there is up-regulation of caspase −3 activities early (at one day) in the KA induced excitotoxic group compared to the control group. This observation is in accordance with results of previous studies showing that the content of caspase-3 mRNA in the spinal cord tissue was raised after KA injury ***(Nottingham and Springer 2003 and Kuzhandaivel et al. 2010).*** In the present study the down-regulation of caspase-3 expression following estrogen treatment indicated the neuroprotective effect of estrogen. This finding is in harmony with results published with ***Cuzzocrea et al (2008) and Kachadroka et al (2010)***.

#### Astrocytes, microglia and IL-1β expressions

The neurotoxic action of kainate is associated with activation of microglia and astrocytes, which proliferate and migrate into the affected region ***(Somera-Molina et al., 2007 and Cho et al., 2008)***. IL-1β, a proinflammatory cytokine, was produced abundantly by stimulated astrocytes and microglia cells ***(Colton and Wilcock, 2010)***. In our study, the expression of mRNA for IL-1β was significantly increased after KA injection. Our results are also those obtained by previous investigators, who stated that primary cultures of microglia were significantly stimulated by KA, and showed increased IL-1*β* levels **(*Zheng et al., 2009 and Jarvelä, 2011).***

After activation, astrocytes secrete several neurotoxic substances and express an enhanced level of glial GFAP. The increase in GFAP production was considered a hallmark of astrogliosis and an early biomarker of neurotoxicity ***(Liu et al., 2003).*** The up-regulation of GFAP was confirmed in the current study. These findings are in agreement with previous studies that have noted steady increases in astrocyte activation and GFAP expression up-regulated following SCI ***(Chen et al., 2005 and Mitra et al., 2013)***. On the contrary, the results previously published by ***Ding et al. (2000)*** showed that the levels of hippocampal GFAP were unaffected at one day of injury and a 20% decrease was noticed in the amygdala/pyriform cortex after systemic administration of KA. Two days later, GFAP increased similarly in both regions and continued to one month. This result could be attributed to the use of systemic administration of kA not stimulating GFAP expression as rapidly as intra-spinal injection.

The activation of nuclear factor kappa B (NFκB), a pro-inflammatory transcription factor, is an essential step in the progress of inflammation. Estrogen treatment has been found to block activation of NF-κB in acute SCI by interfering with the translocation of NFκB, from the cytoplasm to the nucleus ***(Wu et al., 2013)***. In the present study the down-regulation of IL-1β and Iba-1 in the estrogen treated group compared to the KA group might indicate the neuroprotective effect of estrogen. These findings are closely comparable to those of previous studies which revealed that the expression of several cytokines (including interleukin IL-1β, IL-6 and tumor necrosis factor alpha (TNF-œ)) was decreased at the injury area after 17b-estradiol injection pre- and post-SCI ***(Samantaray et al., 2011 and Siriphorn et al., 2012).***

Astrocytes represent main target cells for estrogen in the CNS. The astrocyte also has the greatest potential to induce the neuroprotective effect of estrogen (***Dhandapani and Brann, 2003)***. Our approach revealed that the expression of GFAP after estrogen therapy was variable. At the 1st day post estrogen treatment, the expression of GFAP was markedly up-regulated compared to the KA excitotoxic group, but after that the expression was significantly decreased to levels nearly indistinguishable from control group. This result is in harmony with results of previous studies which revealed that a high dose of 17β-estradiol delivered immediately in rat after SCI and continued for 21 days, enhances the immunoreactivity of GFAP and vimentin followed by decrease of this immunoreactivity at 50 days after injury ***(Ritz and Hausmann, 2008).*** This dramatic increase of GFAP on the 1st day is attributed to the increase of astrocyte numbers as the main target for estrogen to induce the neuroprotective effect and attenuate the inflammatory response after KA injury. The resolution of inflammation after that, resulting in decreases in the number of astrocytes was accompanied by steady attenuation of GFAP expression until it reached levels nearly indistinguishable from the control group.

### Therapeutic time window for stem cell transplantation into injured spinal cord

***Ogawa et al. (2002)*** mentioned that adult spinal cords are not completely non neurogenic for transplanted NSCs. Rather, a brief therapeutic time window can permit successful implantation. This narrow window may change since the host spinal cord’s microenvironment quickly changes following injury.

But the chronic phase of SCI isn’t likely to be suitable for stem cell implantation because of lack of inducing factors for neurogenesis, glial scarring, or enlarged cyst formation, any or all of which might inhibit the survival of engrafted cells and neuronal regeneration ***(Okano, 2002)***. Most of transplantation studies after SCI are done within 1–2weeks post injury when the potential for cell replacement is expected to be optimum ***(Okano, 2002)***. Based on this we have transplanted the h-GFP-OBNSCs at 1week (sub-acute phase) after KA induced excitotoxic injury. ***Karimi et al. (2006)*** also confirmed that the sub-acute phase permits the survival of stem cells. They demonstrated great success in optimizing the survival of transplanted adult neural precursor cells 2 weeks (sub-acute phase) after spinal cord injury. On the other hand they only detected dead grafted cells after 8 weeks of injury (chronic phase).

### Immuno-histochemical assessment of hGFP-OBNSCs

Long-term survival of engrafted cell populations is very important to researche that demonstrates functional recovery correlated with cell replacement and/or incorporation within the host circuitry, as in the present study. Previous studies revealed that successful engraftment in transplantation strategies of the CNS is very difficult to achieve, particularly for xenografts. Moreover, many studies failed to carry out suitable control of host immunorejection, making the evaluation of these parameters hard to do ***(Salazar et al., 2010).*** In that regard the modulation of an immune response with immune-suppressant medications (such as tacrolimus or cyclosporine, used in the current study) was critical to avoid host-mediated rejection against h-GFP-OBNSCS xenografts ***(Hooshmand et al., 2009)***. The present work exhibited that the h-GFP-OBNSCs were able to survive for 6 weeks after subacute transplantation into injured spinal cord.

Furthermore, no tumors or signs of immune rejection were recorded during the current study. Engrafted cells not only migrated for considerable distances in the spinal cord, they also showed a tropism toward the lesion site. This result agrees with our previous results revealing that all transplanted rats exhibited successful engraftment with h-NGF-GFP-OBNSCs after more than 9 weeks without tumor formation or abnormal morphology in the spinal cord (***Marei et al., 2016***). Similar results were also obtained after sub-acute transplantation of pre-labeled human embryonic stem cell-derived oligodendrocyte progenitor cells (hESC-derived OPCs) by ***Keirstead et al (2005)***, adult neural precursor cells by ***Karimi et al (2006)*** and human NSCs by ***Salazar et al (2010)*** in rat spinal cords. Conversely, the results of other previous studies have shown that a small number of green-fluorescent bone marrow mesenchymal stromal cells (GFP-transgenic BMSCs) were found within the rats’ spinal cord at 2 days, but none at 7 days post-initial transplantation (***Vanecek et al., 2012; Nakamura et al., 2013 and Urdzíková et al., 2014)***. These results confirm that MSCs have no affinity to be integrated into the spinal cord.

Quantitative analysis of OBNSCs showed that at the 6^th^ week after implantation, the average total number of GFP-OBNSCs found in the spinal cord was 380470 of the original number (160,000) that were transplanted. These values represent an approximately 2.38 fold increase in the initial cell population transplanted. 40.91% showed immune reactivity for mature neurons, 36.36% showed immune reactivity of astrocytes and 22.73% showed immune reactivity of oligodendrocytes.

The fate of any engrafted cells and the capacity of these cells for neurorepair is influenced by the niche into which the cells are implanted. This niche could reverse the intrinsic properties of engrafted cells and affect mechanisms of recovery. These results indicate that the niche in this study directs the differentiation of h-OBNSCs mainly towards motor neurons rather than oligodendrocytes and astrocytes, to replace the neurons in this area damaged by KA injection. Conversely, the niche following the mechanical ***(Marei et al., 2016)*** and contusive models ***(Hooshmand et al., 2009)*** direct differentiation mainly towards astrocytes and oligodendrocytes rather than neurons. This is attributed to the idea that mechanical and contusive injuries are accompanied by a prolonged secondary inflammatory response, especially in white matter; and that large caliber axons are preferentially targets of mechanical injury ***(Blight and Decrescito, 1986)***.

A similar sequence of events has been stated by ***Jin et al. (2014)*** who established that implanted murine induced neural stem cells (iNSCs) in the spinal cord mainly differentiated into neurons (53.63 ± 0.32%), astrocytes (15.05 ± 0.31%), and oligodendrocytes (4.12 ± 0.23%) at 3 months after implantation. On the contrary, many other studies reported a predominant differentiation of transplanted cells towards astrocytes after implantation of NSCs into spinal cord ***(Pallini et al., 2005; Macias et al., 2006 and Marei et al., 2016).***

### Motor activity Assessment

The locomotor deficits seen following KA injection are attributed to degeneration of primary neurons of the hind limb central pattern Generator for locomotion (CPG) that exists in the rostral lumbar enlargement (between T13–L2). Therefore, damage to this enlargement can lead to severe deficits in locomotor performance by removing important components of the CPG ***(Cazalets et al., 1994, 1995; Cowley and Schmidt. 1997)***. The CPG is certainly critical to hind limb motor function ***(Cowley and Schmidt, 1997)***. Replacement of the CPG must be considered paramount if recovery in locomotor function following SCI is the goal ***(Magnuson and Trinder, 1997)***.

The current study revealed a noticeable improvement in locomotor functions both in estrogen and hOBNSCs+estrogen groups compared to the KA injured animals, but this improvement was faster and more apparent in the hOBNSCs+estrogen group. The results of the histological and immunohistochemical analyses might indicate that this apparent and faster recovery could be due partly to the neuroprotective effect of estrogen helping to alleviate tissue loss from secondary injury. This would be expected to provide a permissive environment suitable for cellular ingrowth. This environment also enhances the grafted cells to integrate into the host tissue and replace damaged or lost cells. Our findings are substantiated by the conclusion of ***Lee et al (2012)*** who has reported that male rats that received 17β-estradiol at 7 and 24 h after SCI had higher BBB scores and performed better in the footprint analysis, grid walk test and inclined plane test at 3–4 weeks after-injury. Our findings are in contrast to the results obtained by ***Marei et al (2016)*** who demonstrated that there was no improvement in locomotor activity in hNGF-GFP-OBNSCs transplanted rats at any time point of the study. He also asserted that the severity of injury and the presence of extensive gliosis may be the underlying reason that led to decreased ability of implanted human OBNSC to restore the damaged nerve tracts, and gray mater associated with contusion SCI.

Collectively, these data suggest that the combination of estrogen and hOBNSCs promotes a compensatory mechanism to reorganize neural networks in the central pattern generator for locomotion by generating mature neurons, oligodendrocytes and astrocytes.

